# A Machine Learning–3D Microvessel Platform Identifies Kinase Targets Restoring Blood-Brain-Barrier Endothelial Integrity

**DOI:** 10.64898/2025.12.11.692686

**Authors:** Yu Jung Shin, Ling Wei, Sumin Hong, Christina Ren, Micah Jeffrey, Jessica Cardenas, Harry Higgins, Nhi N. T. Ho, Kira Evitts, Joseph D. Smith, Jing-fei Dong, Alexis Kaushansky, Ying Zheng

**Author notes:** co-corresponding authors: Ying Zheng; Alexis Kaushansky. **Competing interests:** The authors have declared that no conflict of interest exists.

## Abstract

Disruption of the brain endothelial barrier is a hallmark of traumatic brain injury (TBI), and contributes to cerebral edema, coagulopathy, and delayed neurological deficits, yet there are no mechanism-guided therapies that directly stabilize human brain vessels. Here we used a three-dimensional (3D) human brain endothelial vessel platform with quantifiable macromolecular barrier function to assess responses to thrombin and plasma from patients with TBI. This endothelial-focused 3D system isolates the central cellular regulator of blood–brain barrier function and enables mechanistic analysis of endothelial signaling and drug responses as a foundation for personalized vascular therapeutics. By integrating this platform with machine learning-guided kinase analysis, we identified kinase pathways associated with barrier stabilization versus disruption and nominated inhibitors that reversed thrombin- and TBI plasma–induced leakage as well as compounds that exacerbated endothelial loss. This work establishes an important first step toward precision vascular therapeutics in TBI, by linking patient-derived plasma phenotypes to specific endothelial barrier responses and pharmacologically tractable kinase signaling, and it provides a generalizable framework for mechanistic drug discovery across neurovascular disorders.

**SIGNIFICANCE STATEMENT:** Blood-brain barrier (BBB) disruption drives cerebral edema, coagulopathy, ischemia, and neurological deficits after TBI. Targeting kinases as therapeutics is promising but challenged by complex inhibitor profiles and patient variability. We present a machine learning-guided platform built on 3D perfusable human brain endothelial-centric microvessels to systematically screen and select kinase inhibitors that reverse TBI-induced endothelial barrier breakdown. By pinpointing key kinase and gene regulators and demonstrating restoration of endothelial barrier integrity in patient-derived plasma models, our approach provides a transformative strategy for stabilizing the endothelial barrier after TBI and offers a versatile framework for therapeutic discovery across neurovascular disorders.

## INTRODUCTION

Traumatic brain injury (TBI) is marked by both immediate structural damage and delayed secondary cerebral and systemic injuries, including tissue edema, ischemia, and inflammation (1, 2). It presents with highly heterogenous clinical outcomes, spanning a broad spectrum of vascular, neurological, and cognitive impairments (3, 4). A key early pathological event of TBI is the disruption of the blood-brain barrier (BBB), which strongly correlates with poor prognosis (5). The BBB is comprised of brain microvascular endothelial cells (ECs) at the innermost layer, connected by tight and adherens junctions, and perivascular cells, forming a selective barrier that regulates molecular exchange between bloodstream and brain parenchyma (6, 7). The breakdown of endothelial barrier exposes the central nervous system (CNS) to circulating neurotoxic factors in blood, promoting neurodegeneration and systemic complications (8–11). Therefore, understanding mechanisms of brain endothelial barrier regulation is essential for developing targeted therapeutic strategies for TBI and has broad implications for treating other related CNS disorders.

Endothelial barrier integrity is dynamically regulated by kinase signaling pathways that modulate junctional stability, cytoskeletal tension, and focal adhesions (12, 13). Kinases such as Src family kinases (SFKs) exert context-dependent effects: disrupting barriers via phosphorylation and internalization of vascular endothelial (VE)-cadherin during inflammation, while promoting barrier restoration as inflammatory processes subside (13–17). Additionally, SFKs regulate focal adhesion and actin cytoskeletal remodeling, thereby integrating signals from both cell-cell and cell-matrix interactions to barrier function (18). However, the therapeutic targeting of kinases remains challenging due to their temporal dependent roles in endothelial barrier integrity and pleiotropic effects of kinase inhibitors (KIs), which are influenced by timing, biological state and individual patient variability.

Kinase regression (KiR) is a machine learning approach that employs elastic net regularization and residual activity profiles of KIs against hundreds of kinases to infer key kinase regulators underlying specific biological phenotypes (12, 15, 19–21). While this approach has been successfully implemented in 2D endothelial cell cultures to study inflammatory responses and temporal phosphorylation dynamics (12, 15), its application in more physiologically relevant 3D microvessel models remains limited. Compared to a static 2D endothelial monolayer systems, 3D microvessels has less throughput but recapitulate critical *in vivo* features, including perfusable lumens with quiescent endothelium, flow dynamics, and cell-cell and cell-matrix interactions (22–24). Furthermore, the high surface-to-volume ratio enables efficient screening using small volumes of patient-derived plasma samples. Importantly, the 3D microvessel system also provides a robust barrier model in which functional vascular responses to input stimuli can be quantitatively assessed through permeability measurements. Given the inaccessible human brain microcirculation, integrating KiR with 3D microvessels provides a scalable, and cost-effective platform to elucidate kinase-mediated barrier regulation and identify candidate therapeutics that restore barrier integrity.

Here, we combine machine learning-guided KI screening with a 3D perfusable brain endothelial microvessel model and regression-based kinase analysis to identify key regulators of endothelial barrier integrity and screen small molecule KIs for attenuating secondary injury in a TBI plasma challenge model. This integrative, system-level approach advances mechanistic insight into brain endothelial barrier regulation and provides a foundation for the development of targeted, personalized interventions for TBI. Moreover, the framework established in this study is broadly applicable to a range of vascular dysfunction-related diseases.

## RESULTS

### Perfusable 3D human brain endothelial microvessels respond to TBI plasma

To study human brain endothelial barrier integrity, we developed a 3D perfusable microvessel platform using soft-lithography and collagen injection molding, as we previously described (22). The model consists of two hexagonal networks (100 µm diameter) connected by a central straight vessel (150 µm diameter), mimicking the branching architecture of brain vasculature (Fig. **1A**). Each unit features an inlet and an outlet to enable direct seeding of human brain microvascular endothelial cells (HBMECs) and culture under perfusion. By day 7, HBMECs formed fully lumenized microvessels with a robust endothelial barrier verified by VE-cadherin immunostaining (Fig. **1B**). The perfusability and precise geometry allowed quantitative assessment of barrier permeability by measuring the permeation of fluorescein isothiocyanate (FITC) dextran from vessel lumen into the surrounding collagen matrix. Under resting conditions, microvessels demonstrated low permeation to both 70kDa and 500kDa FITC-dextran, indicating intact barrier integrity (Fig. **1C**). Barrier integrity was quantified using a ‘permeation index’, defined as the normalized permeation distance of FITC-dextran after three minutes of perfusion (Supplementary Fig. S1). Ultrastructural transmission electron microscopy (TEM) showed a continuous endothelial lining with intact adherens and tight junctions, abundant vesicles, Weibel-Palade bodies, and mitochondria, all features indicative of a quiescent, functional endothelium at resting states (Fig. **1D**). This 3D BBB model, along with quantitative measurements of barrier function using the smaller-sized 70kDa dextran, was subsequently employed to assess endothelial barrier response to KIs and activation stimuli, including thrombin and peripheral blood plasma from TBI patients.

**Figure 1.**
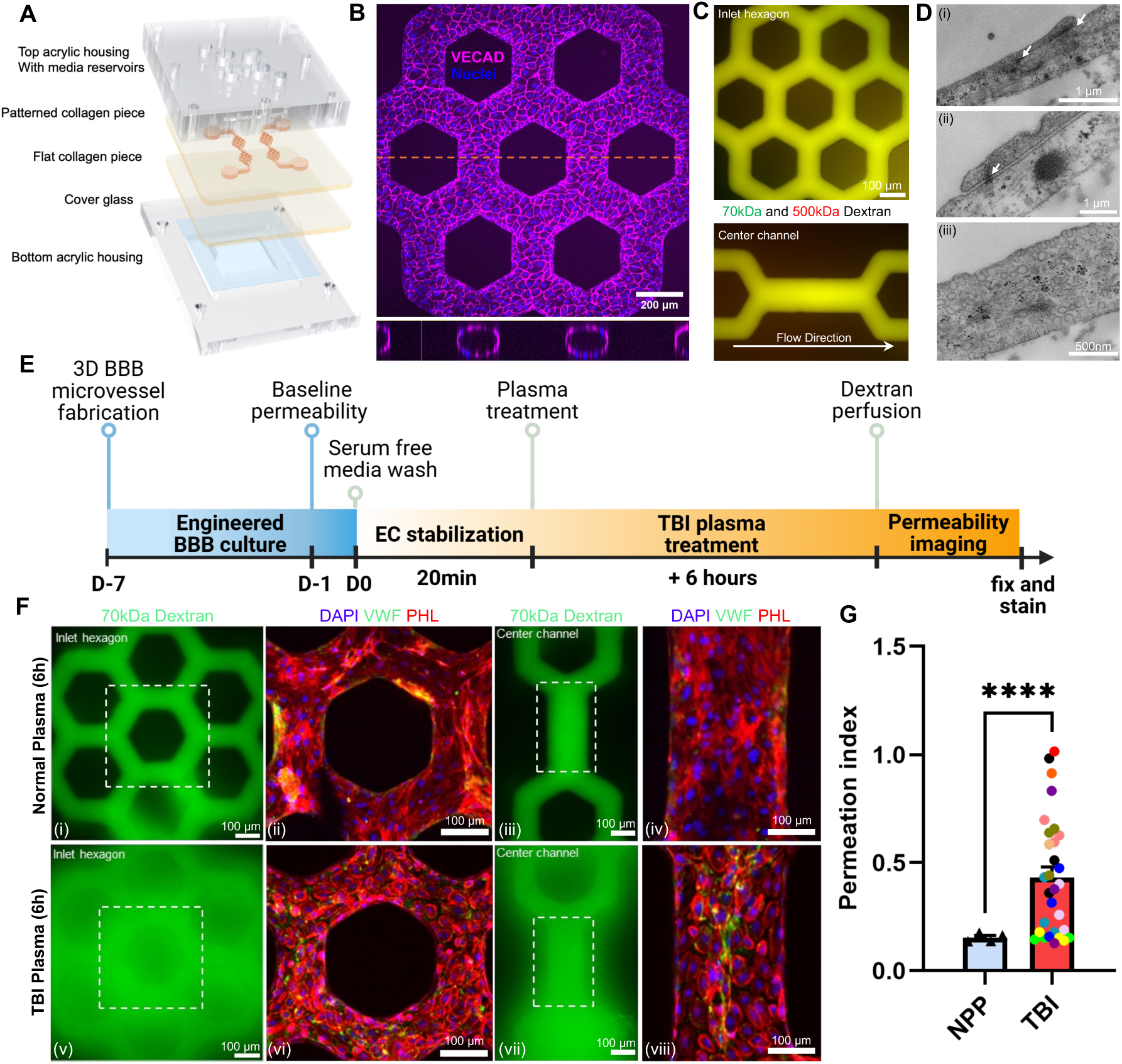
**A 3D perfusable endothelial blood-brain-barrier (BBB) model responds to plasma from traumatic brain injury (TBI) patients.** A. Schematic of the 3D microvascular platform featuring hexagonal network architecture. B. Immunofluorescence images of HBMEC-lined microvessels stained for VE-cadherin (magenta). Orthogonal cross-sectional view shows circular, intact lumens and well-formed junctions after 7 days in culture. C. Representative image of day 7 microvessels perfused with FITC–dextran (70 kDa, green) and TRITC–dextran (500 kDa, red). Merged channels show luminal retention of both tracers, confirming an intact endothelial barrier. D. Transmission electron microscopy (TEM) of resting vessels reveals that HBMECs form (i,ii) well-developed adherens and tight junctions (white arrows) with neighboring cells. Additional features include (iii) abundant vesicle trafficking, indicative of functional endothelium. E. Schematic of the permeability imaging protocol used to assess barrier function following 6-hour treatment with plasma from TBI patients. F. Representative images of 70 kDa FITC-dextran perfusion in 3D BBB microvessels after 6 h exposure to (i,iii) pooled normal plasma and (v,vii) TBI patient plasma. Normal plasma preserved barrier integrity, whereas TBI plasma induced significant leakage. Immunostaining of F-actin (phalloidin; red) and von Willebrand factor (vWF; green) shows endothelial retraction and vWF fiber formation in TBI plasma-treated vessels (vi,viii), which were absent in vessels treated with normal plasma (ii,iv). G. Quantification of the permeation index reveals a three-fold increase in TBI plasma–treated microvessels compared to normal plasma controls (mean ± s.e.m.; N=4 for pooled healthy plasma for normal plasma controls; N =13 biological replicates for TBI plasma, n=1-3 technical replicates per biological condition). Each color represents the same biological TBI plasma samples. ****p < 0.0001; Welch’s t-test.

TBI severity is partly driven by BBB breakdown in response to systemic pro-inflammatory and hypercoagulable signals in plasma. To investigate TBI disease mechanisms, we perfused 3D brain microvessels with plasma from TBI patients and evaluated changes of endothelial barrier integrity. Microvessels with confirmed baseline integrity (D-1), defined by the absence of focal dextran leakage, (Fig. **1E** and Supplementary Fig. S1) were perfused with TBI plasma for 2 h or 6 h. Plasma samples from over twelve patients were perfused through the system, for nine of these patients, at least three microvessels were used as technical replicates, yielding a total of over 40 vessels. After 2h, barrier function did not differ significantly between the healthy donor plasma and TBI plasma conditions (Supplementary Fig. S2). In contrast, a 6 h perfusion with TBI plasma induced a significant increase in 70kDa FITC-dextran leakage (Fig. **1F** v and vii and Supplementary Fig. S2) compared to the healthy donor plasma (Fig. **1F** i and iii and Supplementary Fig. S2). Despite variability between TBI donor plasma, most TBI plasma induced significant barrier disruption (Fig. **1F** and Supplementary Fig. S3). Overall, the TBI patient plasma induced a four-fold increase in permeability compared to healthy pooled plasma (Fig. **1F** and Fig. **1G**).

These findings are consistent with *in vivo* observations from murine TBI models, which reported vascular leakage and hematoma formation within 6 h of fluid percussion injury (25). Post-perfusion analysis in 3D microvessels revealed endothelial cell remodeling and luminal formation of von Willebrand factor (VWF) fibers, indicating endothelial activation and a pro-thrombotic state (Fig. **1F** vi and viii). In contrast, 3D microvessels perfused with healthy pooled plasma remained quiescence with intact barrier (Fig. **1F** ii and iv). These results validate the 3D microvessel model as a physiologically relevant platform for studying TBI-induced vascular dysfunction and hemorrhagic coagulopathy.

### Recapitulating TBI-linked vascular instability with thrombin in 3D microvessels

To understand the procoagulant environment in TBI, we analyzed plasma samples from additional ∼ 30 TBI patients in the cohort and observed significantly elevated levels of phosphatidylserine-positive extracellular vesicles (PS^+^ EVs), known to enhance thrombin generation and promote hypercoagulability (26) (Fig. **2A**). To validate these findings in a controlled setting, we examined the coagulant environment in a mouse fluid percussion injury model. Within 3 h post injury, levels of PS^+^ EVs increased (27) (Fig. **2B**), plasma fibrinogen levels decreased (Fig. **2C**), and thrombin generation was elevated (Fig. **2D**), collectively indicating a hypercoagulable state compared to sham controls.

**Figure 2.**
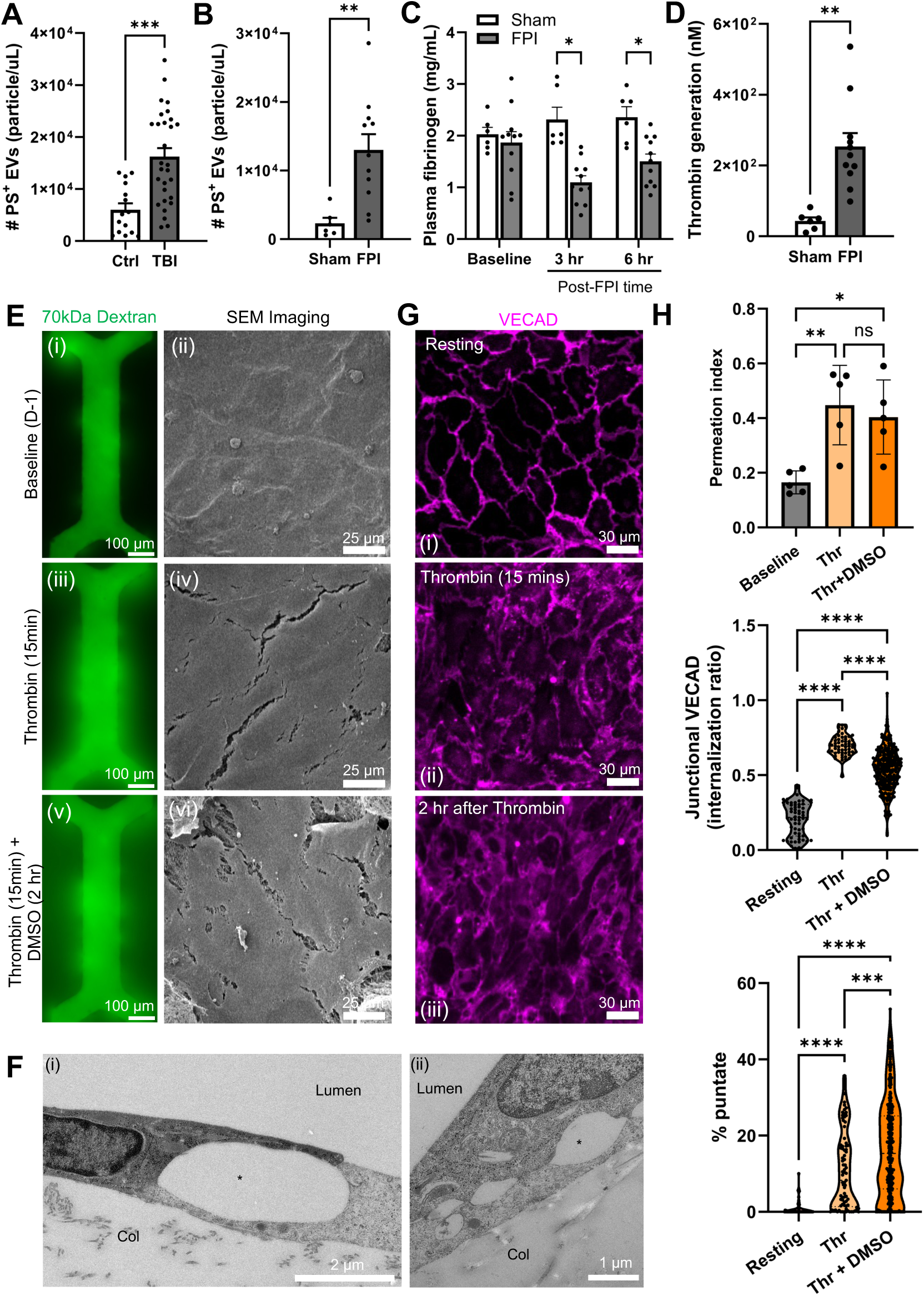
**Thrombin activation in TBI patient plasma induces barrier breakdown in 3D BBB microvessels.** A. Plasma levels of phosphatidylserine-positive extracellular vesicles (PS^+^EVs) in peripheral blood from TBI patients and age- and sex-matched healthy controls (N = 29; Student’s t-test). B-D. Coagulation profile of mice subjected to severe fluid percussion injury (FPI) or sham surgery: B, plasma levels of PS⁺ EVs; C, plasma fibrinogen (a thrombin substrate); and D, thrombin generation (n = 12–16 per group; Student’s t-test). E. Permeability and scanning electron microscopy (SEM) analysis of 3D brain microvessels following stimulation with 1 U/mL thrombin. (n = 5; one-way ANOVA with Tukey’s post hoc test.) F. TEM confirms thrombin-induced damage with discontinuous junctional contacts and gaps formation (marked with *). G. Representative immunofluorescence images of VE-cadherin at (i) baseline, (ii) 15 min after thrombin exposure, and (iii) 2 h after DMSO treatment following 15 min thrombin exposure. H. Quantification of permeation index showed thrombin treatment significantly increased permeability after 15 min; barrier function remained impaired 2 h later relative to baseline (upper panel). Quantification of endothelial remodeling showed increased VE-cadherin internalization ratio (middle panel); and percentage of punctate VE-cadherin junctions (lower panel) after thrombin treatment. Data is shown as mean ± s.e.m. *p < 0.05, **p < 0.01, ***p < 0.005, ****p < 0.001; one-way ANOVA with Tukey’s post hoc test.

Given the significantly increased thrombin generation and its central role in TBI-associated vascular dysfunction (27–29), we examined its direct impact by perfusing microvessels with 1 U/mL of thrombin and assessed permeability and junctional integrity at 15 min and 2 h post-treatment (Fig. **2E-H**). Under baseline conditions, microvessels showed minimal dextran leakage into the collagen matrix, associated with an intact and smooth vessel lumen surface by SEM imaging and continuous VE-cadherin circumferential staining (Fig. **2E** i-ii, and Fig. **2G** i). Thrombin rapidly induced dextran leakage, elevated permeability, long intercellular junctional gaps, and substantial remodeling of VE-cadherin by 15 min (Fig. **2E** iii-iv and Fig. **2G** ii). At 2 h post-thrombin exposure, the junctional gap openings were incompletely repaired (Fig. **2E** v-vi); gaps remain at cell-cell junctions (Fig. **2F**) and the VE-cadherin junctions have not completely reformed (Fig. **2G** iii). Quantification confirmed sustained barrier breakdown persisted throughout the 2 h observation window following a 15 min thrombin challenge (Fig. **2H**, upper panel). VE-cadherin became punctate and relocalized to intracellular compartments (Fig. **2H**, lower two panels, and Supplementary Fig. S4), while cells adopted elongated with reduced circularity and roundness (Supplementary Fig. S4). These findings confirm thrombin as a potent acute disruptor of the brain endothelial barrier and highlight the utility of this 3D microvessel model for dissecting TBI-induced coagulopathy and endothelial dysfunction.

### Machine learning-based KI panel on endothelial barrier regulation in a 3D microvessel model

We previously screened 28 KIs in 2D brain endothelial monolayers to identify regulators of resting and thrombin-induced barrier disruption (15). To translate the higher-througput KiR framework from 2D culture to a lower-throughput 3D brain microvessel platform, we first applied an elastic-net regression approach to our previously published 2D HBMEC permeability dataset (15). By training models over 100 iterations on randomly drawn subsets of 5 to 25 KIs (α = 0.8, condition-specific cross-validation) and evaluating median absolute percent deviation, we found that panels of 10 KIs consistently achieved the lowest prediction error (**Fig. 3A** and **3B**). We therefore selected the top-performing 10 compounds from these iterations and supplemented them with two clinically approved BCR-ABL inhibitors, Bosutinib and Dasatinib, which have divergent endothelial effects in 2D models. The final 12 KIs, listed in Supplementary Table S1, with previously characterized by kinase residual activity (21), exhibited diverse and partially overlapping targets across the kinome (Fig. **3C**). Staurosporine, K-252a (a staurosporine analog), and SB218078 exhibited the broadest inhibitory profiles, with K-252a and Staurosporine each inhibiting over 200 out of 291 screened kinases. These broad activities suggest a need for a system-level approach to dissect kinase-mediated barrier-regulatory pathways.

**Figure 3.**
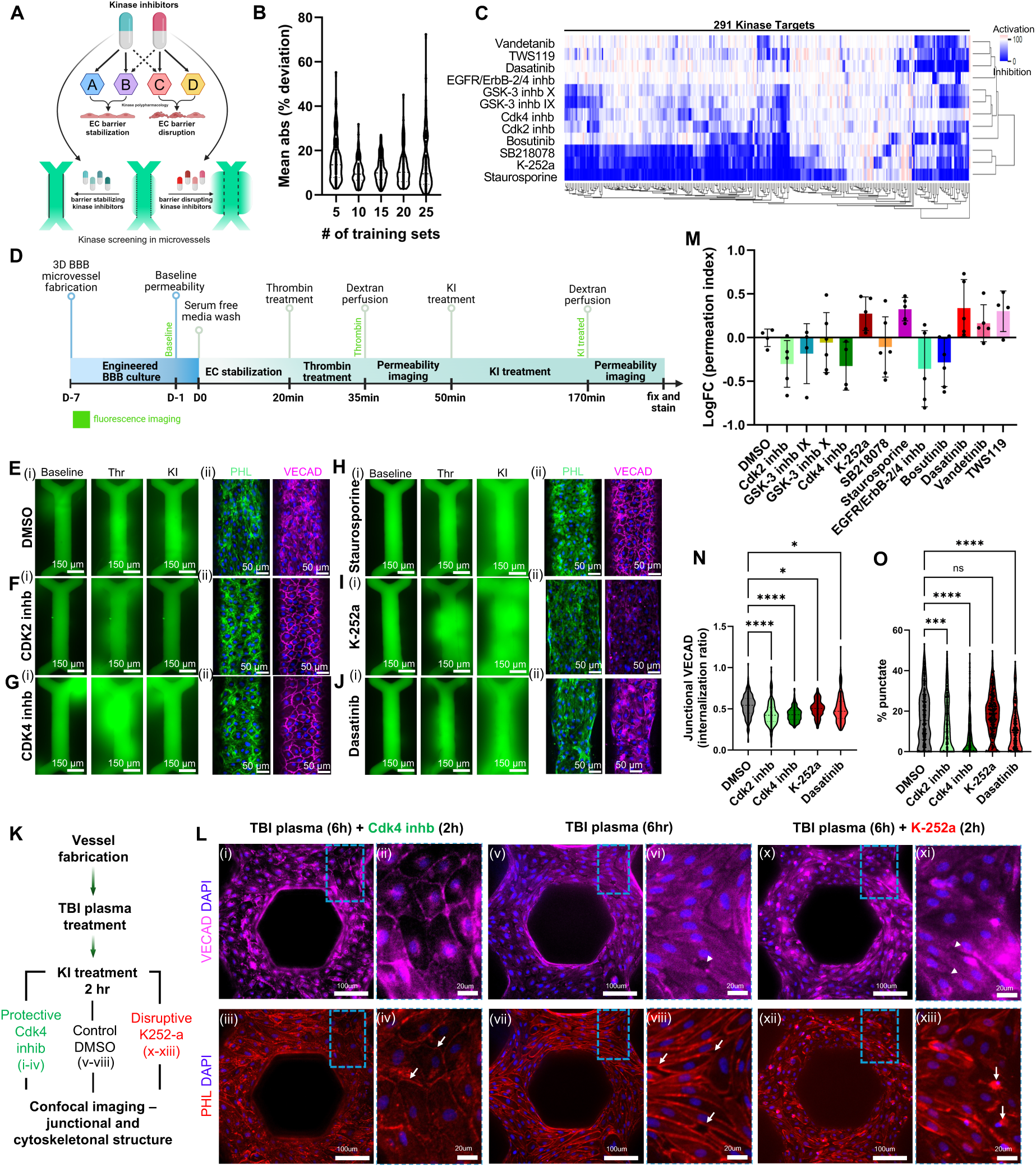
**Machine-learning guided selection of KI panels differentially modulate thrombin-induced barrier disruption in 3D BBB microvessels.** A. Schematic of KI screening workflow using the 3D BBB microvessel for permeability measurement. Schematic produced with Biorender.com B. *in-silico* elastic-net regression analysis identifying ten KIs with lowest prediction error, based on median absolute percent deviation in 28 previously screened KIs in 2D monolayer. C. Final KI panel selection for 3D screening, including 10 from training model iteration in (B) and 2 FDA approved BCR-ABL inhibitors (Bostutinib and Dasatinib; underlined) with known endothelial effects, showed a diverse residual activity for 291 kinase targets. Two-way hierarchical clustering based on normalized residual activity (relative to un-treated controls = 100); values <100 indicate inhibition, >100 indicate activation. D. Schematic of the permeability imaging workflow used to assess the effect of KIs following thrombin exposure. Permeability imaging was conducted in two phases: immediately after thrombin treatment (15 min) and after subsequent KI exposure (2 h), enabling direct within-vessel comparison of barrier disruption and recovery. E-J. (i) Representative images of 70 kDa FITC-dextran perfusion in 3D microvessels at baseline (day –1), after thrombin activation (15 min), and after 2 h of treatment, and (ii) Immunofluorescence images of 3D microvessels fixed immediately after permeability imaging, showing VE-cadherin (magenta) and DAPI (blue) and actin (phalloidin, green). The conditions contain (E) DMSO control; (F) Cdk2 inhibitor; (G) Cdk4 inhibitor; (H) staurosporine; (I) K-252a; and (J) dasatinib. K. Schematic of the permeability imaging workflow. Microvessels were treated with TBI patient plasma for 6 h, followed by 2 h treatment with either previously screened (Cdk2 and Cdk4 inhibitors) or KiR-inferred (ROCK inhibitor and AG490) barrier-restorative KIs. L. Immunofluorescence images of 3D microvessels following TBI plasma perfusion: (i-iv) Cdk4 inhibitor shows partial recovery of VE-cadherin (VECAD; magenta) at cell junctions (i,ii; arrowheads) and actin cytoskeleton reorganization (iii,iv; arrows). (v-viii) TBI plasma alone causes VECAD loss (v,vi) and endothelial retraction (vii,viii; arrows). K-252a treatment exacerbates injury, causing further junctional loss (x,xi; arrowheads) and cell death (xii,xiii; arrows). M. Quantification of log fold change in permeability index across all treatment conditions following thrombin injury and KI treatment (mean ± s.e.m. N = 4–6 biological replicates). Fold change in permeation index (KI/T) was obtained by taking dividing permeation index from Thrombin and KI treated microvessels with permeation index obtained after Thrombin treatment alone in same microvessels. N-O. Quantification of (N) VE-cadherin internalization ratio and (O) percentage of punctate junctions for selected KI treatments. Data are presented as mean ± s.e.m. *p < 0.05, **p < 0.01, ***p < 0.005, ****p < 0.001; one-way ANOVA with Dunnett’s or Tukey’s post hoc test.

To evaluate the capacity of KIs to modulate thrombin induced endothelial injury, we applied each of the 12 KIs to human 3D brain microvessels following exposure to 1U/mL thrombin to assess the changes of vessel barrier integrity. Permeability was assessed at three stages: baseline (D-1), 15 minutes post-thrombin exposure, and after 2 h of KI exposure (Fig. **3D**), all in serum free medium conditions. Only microvessels exhibiting intact baseline barrier function were advanced to thrombin treatment. Permeability was quantified by perfusing 70 kDa FITC-dextran, and cellular architecture was assessed by immunofluorescence, SEM, and TEM (Fig. **3E-M**, Supplementary Fig. S5). Among the 12 KIs tested, Cdk2 inhibitor (Fig. **3F** i), Cdk4 inhibitor (Fig. **3G** i), EGFR inhibitor and bosutinib restored barrier integrity and reduced dextran leakage in thrombin-injured microvessels compared with DMSO controls (Fig. **3E** i). In contrast, K-252a, Staurosporine, and Dasatinib further exacerbated barrier disruption (Fig. **3H-J** i) along with TWS119.

Immunofluorescence confocal imaging revealed elongated actin structures (Fig. **3E** ii; left panel) and cytoplasmic VE-cadherin localization (Fig. **3E** ii; right panel) in DMSO vehicle-treated microvessels, i.e. 150 min after 15 mins exposure of 1U/mL thrombin. In contrast, barrier restorative KIs such as Cdk2 and Cdk4 inhibitors promoted cortical actin architecture and continuous, thick VE-cadherin junctions (**Fig. 3F** and **3G** ii), suggesting an improved junction integrity and restored quiescent EC phenotype. Barrier-disruptive KIs elicited distinct patterns of endothelial dysfunction (Fig. **3H–J** ii). Staurosporine induced actin compaction and irregular, tortuous VE-cadherin junctions (Fig. **3H** ii). K-252a caused loss of VE-cadherin and formation of a dense, contracted actin network (Fig. **3I** ii). The leukemia drug Dasatinib led to endothelial detachment from the collagen matrix, localized luminal narrowing, and pronounced VE-cadherin internalization (Fig. **3J** ii), indicative of disrupted focal adhesion to the basement membrane.

Using Cdk4 inhibitor (barrier-strengthening), and Staurosporine (barrier-disruptive) as contrasting examples, we compared their ultrastructural effects against both resting and thrombin-injured baselines via SEM and TEM (Supplementary Fig. S5). In resting microvessels, SEM showed smooth luminal surfaces and tightly apposed EC borders, with continuous tight and adherens and tight junctions shown in TEM (Supplementary Fig. S5 A). Thrombin activation (with DMSO vehicle control for 2h) produces 1-2 µm intercellular gaps, prominent luminal protrusions (SEM), and on TEM, reduced vesicle density, vacuolization, and lysosomal structures, hallmarks of endothelial stress (Supplementary Fig. S5 B). When thrombin-injured vessels were treated with a Cdk4 inhibitor treatment, ultrastructure closely resembled the resting state: SEM images demonstrated restored continuity of the luminal surface and seamless EC borders, while TEM confirms reformed tight and adherens junctions, rich endoplasmic reticulum, minimal pinocytic vesicles and intact nuclei (Supplementary Fig S5 C). In stark contrast, Staurosporine treatment exacerbated damage beyond thrombin phenotype: SEM showed extensive endothelial delamination, surface pore formation, and exposed collagen matrix, and TEM showed loss of cell-cell and cell-matrix contacts, large vacuoles and nuclear condensation consistent with apoptotic breakdown (Supplementary Fig S5 D).

To evaluate the therapeutic potential of screened KIs during early TBI stages, we perfused microvessels with TBI patient plasma for 6h (Fig. **3K**; schematic overview). Treatment with Cdk4 inhibitor, as a barrier-restorative KI, partially restored VE-cadherin localization at junctions (Fig. **3L** i-ii) and reduced cellular gaps (Fig. **3L** iii-iv), compared with DMSO controls (Fig. **3L** v-viii). By contrast, the barrier-disruptive compound K-252a exacerbated endothelial injury, leading to increased cell death, VE-cadherin internalization and cellular retraction (Fig. **3L** x-xiii).

Quantitative analysis confirmed that K-252a, Staurosporine and Dasatinib trended towards increased dextran permeation, and the Cdk4 inhibitor toward reduced leakage versus DMSO control (Fig. **3M**). However, these differences did not reach statistical significance, likely reflecting inter-vessel and cell state variability in permeability measurements. Quantitative analysis of junctional structure and cellular morphology displayed significantly different and divergent effects between barrier-strengthening and disrupting KI treatments (Fig. **3N-O**). Nevertheless, these observed trends suggest the need for a machine-learning framework to predict and prioritize KIs with more robust barrier-modulating functions for future validation.

### KiR identifies kinase regulators and high-efficacy inhibitors for endothelial barrier restoration

To further elucidate key kinase pathways regulating thrombin-induced barrier disruption, we applied KiR, integrating quantitative permeation index, shown in Fig. 3K, from thrombin-activated BBB microvessels treated with 12 KIs, with their biochemical residual activity profile across 291 human kinases shown in Fig. 3C. Elastic net regularization identified 15 kinase targets associated with either barrier disruption or restoration following exposure of 1U/mL thrombin (vertical column, Fig. **4A**), which was used to infer the effects of 166 untested KIs on endothelial barrier function. Hierarchical clustering of residual kinase activities revealed distinct patterns across the ∼20 top-ranked barrier-restorative (right side, Fig. **4A**) and barrier-disruptive KIs respectively (left side, Fig. **4A**). Inhibition of kinases such as RIPK2, PDGFRB, BLK, FYN, EPHA1, and TXK were associated with increased barrier permeability. Notably, thrombin dose-dependent differences at 0.2 versus 1 U/mL suggested context-specific kinase activity linked to coagulation state (Supplementary Fig. S6). Importantly, regression across both doses converged on seven shared kinase targets, including RIPK2, VRK1, EPHA1, MAP3K14, NEK7, CSNK1G1, and ACVR1B, suggesting core regulators of barrier dysfunction under procoagulant conditions (Fig. **4** **A** and **B,** Supplementary Fig. S7). The corresponding predicted inhibitor panels overlapped by 20 compounds, including TGFβ_RI_inhibitor_III, AG_490, ROCK_inhibitor, p38_MAPK_inhibitor, Rho_K_inhibitor_III, JAK_inhibitor, PKR_inhibitor, and GSK3_inhibitor, suggesting a versatile set of drug candidates. While individual responses may vary, the shared activity profiles offer a rationale basis for developing kinase-targeted therapies capable of stabilizing the endothelium across diverse procoagulant states across patients.

**Figure 4.**
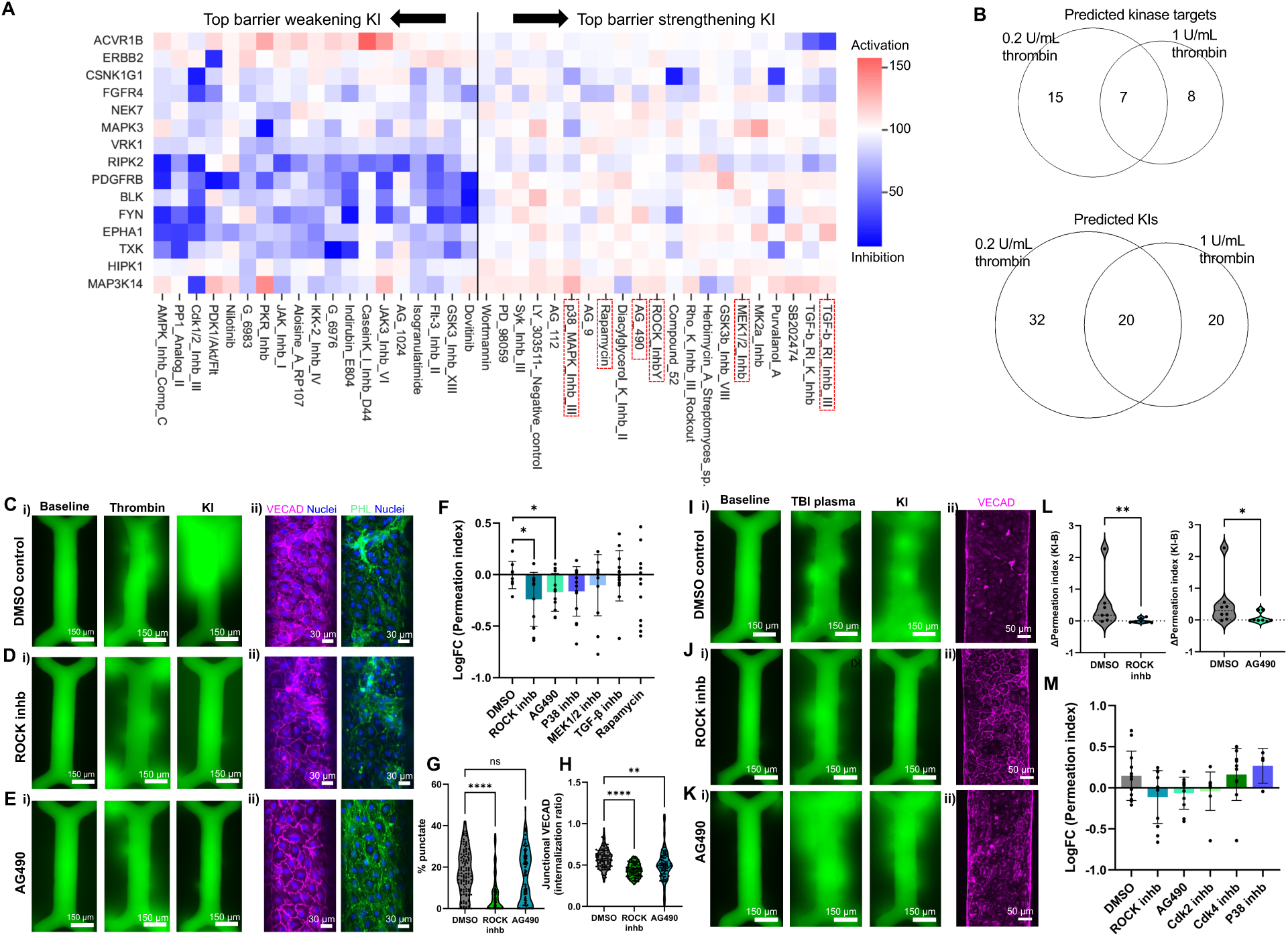
**Kinase regression (KiR) infers new barrier-restorative targets following thrombin-induced BBB disruption and restore barrier integrity in TBI plasma-treated 3D BBB microvessels.** A. Heatmap showing association between kinase targets (y-axis) and kinase inhibitors (x-axis) based on permeability outcomes under 1 U/mL thrombin conditions. Barrier-restorative predictions appear to the right of the vertical axis; disruptive predictions to the left. Red indicates activation; blue indicates inhibition. Red dashed lines indicate selected six KIs for validation. B. Venn Diagrams of predicted kinase targets and alterative KIs for 0.2 and 1 U/mL thrombin treatment, suggesting common and dose dependent injury mechanisms. C–E. Validation of top-ranked new KIs in thrombin (1 U/mL)-treated BBB microvessels. Compared to (C) vehicle control, two KIs—(D) ROCK inhibitor, and (E) AG490 —show the most promising potential for restoring the barrier following thrombin activation. i). permeability measurements, and ii) immunofluorescence images. Right images show nuclei (DAPI; blue) and actin (PHL; green); left: DAPI (blue) and VE-cadherin (magenta). F. Log fold change of permeation index across 6 KiR identified KIs, (N=8–14 vessel replicates for each KI treatment, *p<0.05 Student’s t-test). Fold change in permeation index (KI/T) was obtained by taking dividing permeation index from thrombin and KI treated microvessels with permeation index obtained after Thrombin treatment alone in same microvessels. G-H. Quantification of VE-cadherin structures in microvessels treated with barrier-restorative KIs. (G) % punctate and (H) internationalization ratio. Both parameters were significantly reduced in Rock inhb treatment relative to DMSO control, suggesting stabilization of junctional localization. I-M. Selected KIs restore barrier integrity in TBI plasma-treated 3D BBB microvessels: I-K. (i) Permeability images showing 70 kDa FITC-dextran perfusion (at baseline (day –1); middle: post–6 h TBI plasma treatment; bottom: post–2 h KI treatment) and (ii) VE-Cadherin junctional image immediately after permeability measurement in vessels treated with: (I) DMSO, (J) ROCK inhibitor, and (K) AG490.. L. Change in permeation index after 2 h KI treatment with: (left), ROCK inhibitor; (right) AG490; Changes were quantified by measuring the difference in permeation index obtained after 6hr treatment with TBI plasma to baseline permeation index. (N=7–8 TBI plasma replicates for each KI treatment, *p<0.05, **p < 0.01) M. Log fold change of permeation index for different selected KIs, identifying ROCK inhibitor as the most effective barrier-restorative KI. (N=10–12 vessel replicates for each KI treatment). Fold change in permeation index (KI/TBI) was obtained by taking dividing permeation index from TBI plasma and KI treated microvessels with permeation index obtained after TBI plasma treatment in the same microvessels. Quantification data are presented as mean ±. s.e.m.; *p < 0.05, **p < 0.01, ***p < 0.005 and ****p < 0.001 determined by one-way ANOVA and post-hoc Tukey’s test.

From these overlapping, top-ranked hits, we chose six barrier-restorative KIs, each exhibiting a distinct target profile (Fig **4**A; red boxes), including a ROCK inhibitor, AG490 (JAK2 and EGFR inhibitor), p38 inhibitor, MEK1/2 inhibitor (SL327), Rapamycin (mTOR inhibitor), and a TGF-β inhibitor, for testing in thrombin-injured microvessels. Compared to vehicle (DMSO)-treated controls (Fig. **4C** i), the ROCK inhibitor (Fig. **4D** i), AG490 (Fig. **4E** i), and the p38 inhibitor (Supplementary Fig. S8) significantly reduced thrombin-induced leakage. Quantitative analysis confirmed that AG490 and the ROCK inhibitor provided the most robust barrier-restorative effects (Fig. **4F**), whereas MEK1/2 inhibitor and TGF-β inhibitor had mixed outcomes (Supplementary Fig. S8). These findings support the proof-of-concept value of KiR model in identifying effective barrier-modulating KIs (Fig. **4C-F** and Supplementary Fig. S8). Confocal imaging showed that treatment with ROCK inhibitor or AG490 restored VE-cadherin localization at cell-cell junctions and reduced junctional discontinuities compared with DMSO-treated microvessels, which displayed VE-cadherin internalization and elongated actin fibers (Fig. **4C–E** ii). Quantitative analysis confirmed these effects (Fig. **4G-H**), with the ROCK inhibitor achieving the most substantial junctional recovery (Fig. **4G**). These findings demonstrate the utility of KiR in identifying both mechanistic regulators and candidate therapeutics for restoring endothelial barrier function following thrombin-induced BBB injury.

### KiR-inferred KIs restore barrier integrity in 3D brain endothelial microvessels exposed to TBI plasma

Building on KiR model for thrombin-activated microvessels, we next tested whether KiR-identified KIs could similarly restore barrier functions in TBI plasma-exposed microvessels (N=9 donors; Supplementary Fig. S9). Following plasma perfusion, microvessels were treated with DMSO (control), two KiR-inferred KIs (ROCK inhibitor and AG490), or two previously screened barrier-restorative inhibitors (Cdk2 and Cdk4 inhibitors). As expected, TBI plasma markedly increased permeability compared to baseline (Fig. **4I-K**). DMSO-treated control exhibited sustained leakage, whereas both ROCK inhibitor and AG490 significantly reduced dextran permeation from the microvessels (**Fig. 4J** and **4K**; right panels). Immunofluorescence imaging further revealed that TBI plasma induced VE-cadherin internalization in DMSO-treated vessels (Fig. **4I**), while ROCK inhibitor and AG490 both restored continuous, junctional VE-cadherin localization (Fig. **4J-K** ii). Permeability index quantification corroborated these findings: ROCK inhibitor and AG490 produced the strongest, statistically significant reduction in barrier leakage, while Cdk4 inhibitor conferred only modest protection (Fig. **4L-M**). The advanced barrier-stabilizing effects of these KiR-inferred candidates, surpassing even the initial CDK2/4 hits, highlight the power of machine-learning guided regression to promote therapeutic targeting for endothelial repair.

## DISCUSSION

Kinase signaling pathways play a central role in regulating endothelial barrier integrity by modulating cell-cell and cell-matrix junctions, actin organization, and adherens junction recycling (30–36). In this study, we combined a kinase regression-guided screening approach with 3D perfusable human brain endothelial microvascular models to identify kinases and KIs that regulate vascular permeability under TBI-mimicking conditions. This integrated strategy establishes a proof-of-concept framework for optimizing kinase-targeted therapeutics in treating injury-induced vascular dysfunction. Given that human ECs harbor over 2,000 thrombin-regulated phosphorylation sites (37), kinase-targeting strategies offer a rational path to stabilize endothelial barriers in trauma-induced coagulopathy and secondary neurovascular injury.

The engineered brain microvessels, composed of HBMECs and characterized by high surface area-to-volume ratios, displayed robust junctional architecture and low baseline permeability, while responding sensitively to thrombin and TBI patient plasma. Importantly, the platform required only minimal plasma volumes, making it suitable for screening of multiple KIs or drug candidates per patient sample using limited volumes, addressing a key challenge in translational TBI research where clinical specimens are scarce. By leveraging elastic net regression to our previous 2D permeability screen, we distilled an optimal panel of 10 KIs that minimized prediction error across repeated model training (15), then augmented the core set with two clinically approved BCR-ABL inhibitors, yielding a focused 12-KI set that retained predictive fidelity when transferred into our 3D microvessel models.

### Mechanistic insights from CDK, ROCK, and JAK2 inhibition

Our initial screening panel in 3D human brain microvessels identified inhibitors of CDK2 and CDK4 as barrier-protective candidates under thrombin challenge, despite their canonical association of cyclin-dependent kinases with cell-cycle control (38, 39) rather than acute barrier regulation. The rapid restoration of barrier integrity within hours of thrombin exposure, along with re-established cortical actin architecture and continuous VE-cadherin junctions, pointed to noncanonical CDK2/4 roles in modulating endothelial cytoskeletal and junctional substrates during acute vascular injury. These kinetics suggest that CDK2/4 inhibitors could be deployed in a narrow therapeutic window after TBI to prevent secondary barrier breakdown, with potential downstream benefits in mitigating cerebral edema, coagulopathy, and ischemic injury.

Furthermore, our machine-learning–guided KIR analysis identified alternative kinase targets, including a ROCK inhibitor and the JAK2 inhibitor, AG490 as robust barrier stabilizer. Activation of ROCK pathway promotes endothelial hyperpermeability via RhoA-dependent myosin light chain phosphorylation, promoting actomyosin contraction and junctional tension (40); inhibition of ROCK has been shown to attenuate inflammatory and LPS-induced barrier disruption in multiple vascular beds. Consistent with these mechanisms, ROCK inhibition in 3D microvessels yielded pronounced reductions in FITC-dextran leakage and more complete VE-cadherin relocalization than CDK inhibitors. AG490 likely preserves junctional integrity by attenuating cytokine-induced JAK-STAT signaling and junction protein loss, as reported in post-stroke and IL-6-challenged endothelial models (41). Together, these data validated both canonical (ROCK, JAK-STAT) and less explored (CDK2/4) kinase targets as convergent, mechanism-based strategies for acute endothelial stabilization following TBI and related neurovascular insults.

### Model limitations and future extensions

Despite its predictive performance, the current KiR framework is constrained by its reliance on a single functional readout, the permeation index, which aggregates multiple underlying structural changes into one scalar metric. Future iterations could integrate image-derived metrics such as VE-cadherin continuity, junctional gap area, and cytoskeletal architecture, to more precisely map kinase-specific phenotypes and improve the resolution and accuracy of KiR-based kinase function analysis. Incorporating temporal information, similar to time-resolved KiR approaches (12), may further dissect early disruptive versus late restorative kinase activities under dynamic inflammatory stimuli.

Our system could be further strengthened by incorporating additional neurovascular unit cell types and expanding the scope of KI screening. This study focuses on the endothelial component of the BBB in isolation, whereas *in vivo* barrier properties emerge from coordinated signals among endothelial cells, pericytes, astrocytes and neurons (42, 43). Incorporating additional neurovascular cell types and relevant extracellular matrix cues would enable interrogation of multicellular crosstalk, including pericyte-derived angiopoietin–Tie signaling and astrocyte-mediated barrier reinforcement, potentially uncovering cell-type–specific kinase dependencies (44–47). Furthermore, variability in barrier disruption kinetics across plasma samples and microvessel replicates suggest underlying heterogeneity in endothelial state; single-cell or spatial transcriptomic profiling of microvessels under different plasma conditions may elucidate distinct endothelial subpopulations and improve model robustness and stratification.

Another limitation lies in the composition of the KI training set: only a subset of the 12 KIs were protective under thrombin challenge and kinases driving barrier disruption exerted stronger phenotypic effects than protective kinases. This imbalance biases the KiR model towards predicting disruptive phenotypes and their associated kinase regulators, potentially underrepresenting protective networks that operate with subtler effects. Expanding the training library to include more barrier-restorative compounds, and incorporating longitudinal patient datasets, will help reduce prediction bias, refine kinase-phenotype mapping, and further enhance its capacity to discover protective pathways across diverse injury contexts.

### Translational implications

The reduced efficacy of several barrier-restorative KIs against TBI plasma-induced injury, compared to thrombin-only exposure, suggests that TBI plasma contains additional thrombin-independent factors, such as cytokines, damage-associated molecular patterns, and complement factors, that drive barrier breakdown through parallel or synergistic mechanisms (5, 48). These injuries may be less responsive to thrombin-centric strategies and suggest the need for combination approaches that concurrently target coagulation, inflammatory, and cytoskeletal pathways to provide durable BBB protection after TBI. Future systematic testing of patient plasma against KI panels in a perfusable human BBB model may help prioritize kinase targets that show consistent barrier stabilization across heterogeneous samples, while also revealing patient-specific vulnerabilities amenable to tailored interventions.

Altogether, our work establishes a robust platform that integrates patient-derived plasma, 3D human brain endothelial microvessels, and kinase-informed machine learning to enable high-content, KI-guided functional screening using small plasma volumes. We show that TBI patient plasma elicits multifactorial endothelial injury that is partially reversed by thrombin-focused kinase inhibition, pointing to additional plasma-borne drivers of barrier failure that remain to be defined. By coupling KiR with clinically relevant readouts of barrier function, this approach provides a scalable path to identify convergent kinase targets with broad therapeutic potential while still accommodating inter-patient variability. The ability to nominate clinically advanced inhibitors, such as BCR-ABL or ROCK inhibitors with established safety profiles, that restore endothelial barrier integrity in this model could accelerate repurposing efforts and de-risk translation by leveraging existing pharmacokinetic and toxicity data. More broadly, expanding this platform into multicellular neurovascular unit and establishing them into early-phase clinical trials, where plasma kinase-signaling signatures could be correlated with ex vivo BBB responses, may inform patient stratification, dose selection, and the development of targeted, ultimately personalized therapeutics to restore BBB integrity in TBI and other acute or chronic neurovascular disorders.

## MATERIALS AND METHODS

Comprehensive methods are provided for all aspects of this study, including TBI patient plasma collection, TBI mouse model, coagulation measurements, and details of the 3D microvessels platform, associated assays and KiR models, as the central approach for this study.

### Sex as a biological variable

Sex was not considered a biological variable in this study. Human samples included both male and female donors.

### HBMEC culture and 3D microvessel fabrication

Primary human brain microvascular endothelial cells (HBMECs; Cell Systems, Cat# ACBRI 376) were expanded in tissue culture flasks coated with Attachment Factor (Cell Systems, Cat# 4Z0-210) using EGM-2MV medium (Lonza). Cells were used between passages 4 and 7. The 3D endothelial BBB microvessels were fabricated using soft lithography and collagen injection molding, as previously reported (Zheng et al., 2012). Briefly, polydimethylsiloxane (PDMS) stamps with a hexagonal network pattern (channel diameter: 100 µL) were used to mold microchannel networks into collagen gels. Type I rat tail collagen (15 mg/mL in 0.1% acetic acid) was diluted to 7.5 mg/mL and neutralized with 1N NaOH, 10x M199 media, and EGM2-MV to create a working solution. The gel was loaded into an acrylic jig and incubated at 37°C to crosslink. Top and bottom collagen layers were assembled, and the assembled jig was perfused with media to initiate flow. HBMECs were seeded into the assembled microvessels by perfusing 10 µL of a concentrated cell suspension (7-10 x 10⁶ cells/mL in EGM-2MV) through the inlet. Devices were incubated at 37°C and 5% CO2 and media were replaced every 12 h to sustain gravity-driven perfusion through the microvessel networks. Microvessels were maintained for 7 days prior to use in permeability and imaging assays.

### Plasma, thrombin and kinase inhibitor (KI) treatment

Permeability measurements were conducted on day 7 (D7) BBB microvessels under resting or injured conditions following treatment with thrombin, TBI patient plasma, healthy control plasma, and/or KIs. Baseline permeability of engineered vessels was measured by perfusing 70 kDa FITC-dextran (100 mg/mL in EGM-2MV growth media) or 500 kDa TRITC-dextran (100 mg/mL in EGM-2MV growth media) under a hydrostatic pressure drop (∼1 mmHg) and imaging leakage across the endothelial barrier.

On the day of the assay, microvessels were equilibrated for 20 minutes with serum-free EGM2-MV media. KIs (Selleck Chemicals) and dimethyl sulfoxide (DMSO) vehicle control solutions were prepared from 1 mM stock KIs and diluted to 1 µM in serum-free EGM2-MV media. For screening under resting conditions, each KI and vehicle control (DMSO) were perfused into microvessels for 2 h, followed by immediate dextran perfusion and imaging to assess permeability. Both leaky and intact microvessels were used to assess the barrier-modulating effects of KIs.

For assays involving barrier disruption, microvessels were first treated with thrombin (0.2 U/mL or 1 U/mL in serum-free EGM2-MV media) for 15 min to induce activation. Alternatively, TBI patient plasma and normal pooled plasma was applied directly to microvessel inlets and perfused for 2 h or 6 h to model patient-specific injury responses. Following perturbation, microvessels were washed with serum-free EGM2-MV media for 15 min to remove residual plasma or thrombin, then treated with KIs for 2 h. Following KI treatment, dextran perfusion was introduced to the lumen immediately to assess the changes of barrier function. KIs predicted by the KiR model was prepared and applied using the same protocols as the initial 12 screened KIs. At the end of each permeability assay, microvessels were washed with PBS, fixed in 3.7% paraformaldehyde (PFA) for 15 min, and stored at 4°C for subsequent immunofluorescent analysis.

### Permeability analysis

Fluorescent images were analyzed in Fiji (ImageJ) to track dextran diffusion across vessel walls. Lumen diameter, region of interest (ROI) position, and fluorescence intensity values were extracted and used in custom MATLAB scripts. Endothelial permeability was quantified by calculating the permeability coefficient (K), derived from dextran diffusivity and intensity decay using time-lapse imaging data.

In addition, a complementary metric, the permeation index, was developed to provide a dimensionless measure of dextran leakage particularly useful in thrombin-injured vessels. In these vessels, high baseline leakage at early time points rendered traditional permeability coefficients less sensitive. The permeation index was defined as the normalized distance traveled by fluorescent dextran beyond the luminal boundary. Specifically, the x-position (Δx) corresponding to two-thirds of the peak fluorescence intensity from the center of the lumen was identified. Values of Δx were calculated for both vessel boundaries, averaged, and normalized by the lumen diameter to yield a dimensionless index. This parameter was used as a phenotypic readout for kinase regression analysis.

### Patient plasma collection and processing

Blood samples (anticoagulant: 0.32% sodium citrate) were collected from 42 patients with isolated TBI upon admission to Memorial Hermann Hospital-Texas Medical Center. Isolated TBI was defined as a head abbreviated injury score of ≥3 and all other body regions scores of <3. Clinical details are presented in Supplementary Table 1. Blood was processed within 2 hours of phlebotomy: first centrifuged at 1,200xg for 10 minutes at room temperature to isolate platelet-poor plasma, then again at 12,000xg for 10 minutes at 4°C to obtain cell-free plasma. Plasma was aliquoted and stored at -80°C. Control plasma samples were collected from 16 age- and gender-matched healthy subjects recruited from the Bloodworks Research Institute donor pool.

### TBI mouse model and plasma generation

C57BL/6J male mice (12-16 weeks, 22-25g; Jackson Laboratory, BarHarbor, ME) were subjected to lateral fluid percussion injury (FPI) to the brain under anesthesia, as we previously described (49, 50). Briefly, a mouse was anesthetized, connected to a small animal ventilator (Harvard Apparatus, Hollistone, MA), and restrained to a mouse surgical platform to surgically expose the skull. A 3-mm-diameter hole was drilled into the skull at 2.0-mm posterior from the bregma and 2.0-mm lateral to the sagittal suture with the dura mater intact to mimic closed TBI. A female Leur-Lok was fixed to the craniotomy site using bone cement. The mouse was allowed to recover from the surgery for 60 minutes in a regular cage and then connected to an FPI device (Custom Design & Fabrication, Richmond, VA) through the fixed Leur-lok and with the head unrestrained to mimic TBI injury in real-life. Saline from a Plexiglas cylindrical reservoir was rapidly injected into the closed cranial cavity at a controlled pressure of 1.96 ± 0.3 atm, which we have previously shown to generate severe TBI (49, 51). Sham controls underwent the same surgery without injury. Blood samples were collected from TBI and sham mice at 3 hours post injury immediately after euthanasia under anesthesia (0.38% sodium citrate as anti-coagulant, final concentration). This mouse study was approved by the Institutional Animal Care and Use Committee of the Bloodworks Research Institute.

### Flow cytometry of extracellular vesicles

To assess the release of brain-derived extracellular vesicles (BDEVs), plasma samples (20 µL) were incubated with APC-conjugated annexin V and either FITC-conjugated anti-neuron-specific enolase (NSE, Abcam) or anti-glial fibrillary acidic protein (GFAP, Abcam) antibodies for 20 minutes at room temperature to detect neuronal and glial EVs, respectively. Samples were analyzed on an LSR II flow cytometer (BD Biosciences) using the EV gate defined with Megamix-Plus FSC/SSC beads (Biocytex). EV counts were normalized using Sphero AccuCount beads (Spherotech). Mouse plasma samples were similarly stained with ABC-annexin V and cell markers, and quantified.

### Coagulation Assays

Two assays were utilized to gauge coagulation dysregulation of TBI mice: plasma fibrinogen levels and the rate of thrombin generation, as we previously described^25,38,39^. Fibrinogen levels were measured using a commercial ELISA assay (Abcam) according to manufacturer’s instruction assays. Thrombin generation was measured using a FluCa Kit on Thrombinoscope BV (Diagnostica Stago, Asnières-sur-Seine, France) according to the manufacturer’s instructions^38^. Briefly, 70 μL of phospholipid-deficient porcine plasma (IMVS), 20 μL of FluCa solution (Diagnostica Stago), and 30 μL of plasma from TBI mice (each sample was pooled from blood of 5 TBI mice) were mixed and immediately analyzed at 37°C on a Thrombinoscope Fluoroskan Ascent® reader (Thermo Lab Systems, Helsinki, Finland) at 390 nm excitation and 460 nm emission. For the control, thrombin generation was induced by either 1 pM of tissue factor or 20μL of the MP-Reagent (Diagnostica Stago, Inc., Parsipanny-Troy Hills, NJ), which contains phosphatidylserine, phosphatidylcholine, and phosphatidylethanolamine at a ratio of 20:60:20, thus providing 4μM of total phospholipids to the reaction mix.

### Immunofluorescence staining and microscopy

Following fixation, vessels were permeabilized and blocked in 2% BSA/0.1% Triton X-100 in PBS. Following the 1-hour blocking step, microvessels were stained with primary antibodies overnight at 4°C. For direct staining, microvessels were incubated with 1:100 APC anti-human VECAD (eBioscience, 17-1449-42) and either Alexa Fluor 488 or Alexa Fluor 594 phalloidin (1:100, Invitrogen, A12379 and A12381 respectively). For additional junctional markers (ZO-1 and Claudin 5), samples were first stained with unconjugated primary antibodies followed flourophore-conjugated secondary antibodies. Nuclei were counterstained overnight at 4°C with 1:200 Hoechst 33342 (Thermo Fisher, CC-H1399). Each antibody incubation was followed by three 20-min washes with DPBS. Samples were stored at 4°C until imaging. Confocal imaging was performed on a Yokogawa W1 spinning disk confocal microscope (Nikon Ti2). Z-stack images (5 µm for 10x, 2 µm for 20x) were used for orthogonal and maximum projection rendering in Fiji (ImageJ).

### VECAD quantification and analysis in 3D microvessel

VECAD junctional organization was analyzed from maximum intensity projection (MIP) images using previously validated image analysis tool, the Junction Analyzer Program (JanaP). MIP images were imported into the program and individual cell boundaries were manually traced to quantify edge-localized VECAD junctions. The percentage of punctate junctions was defined as the proportion of the cell perimeter presenting discontinuous, dot-like VECAD staining. The percentage of discontinuous junctions was defined as the fraction of the cell perimeter containing either punctate or perpendicular (non-continuous) junctional structures. More than 150 ECs were analyzed across 3D microvessels treated with 12 kinase inhibitors in thrombin activated condition. More than 80 cells were analyzed for 6 top-ranked kinase inhibitors predicted from KiR.

### Selection of kinase inhibitors for screens

The screening panel of KIs was derived from a previously published 2D HBMEC permeability dataset in which 28 kinase inhibitors were profiled for 2 h barrier effects (15). We employed an elastic-net regression (α = 0.8) via the glmnet_python package (v2.2.1), performing 100 iterations of model training on random subsets of 5, 10, 15, 20, or 25 inhibitors. In each iteration, normalized permeability readouts were regressed against kinase–compound activity data (21) using a condition-specific cross-validation scheme; model performance was evaluated by median absolute percent deviation on the held-out inhibitors. Subsets of 10 compounds consistently minimized prediction error, so we selected the highest-performing 10-inhibitor set (Supplementary Table S1). To incorporate clinically relevant agents with known, divergent endothelial effects in 2D, we then added the BCR–ABL inhibitors bosutinib and dasatinib, yielding the final 12-compound panel.

### Kinase regression analysis

Kinase regression was performed using an elastic net regularization algorithm as previously described (15). The kinase regression (KiR) approach exploits the polypharmacology of a small panel of kinase inhibitors to identify kinase targets driving specific cellular phenotypes and predict the effects of untested kinase inhibitors based on the weighted contribution of kinase activities to the phenotypes (19, 21). For this study, the phenotypic input to the KiR model was the change in permeability of 3D microvessels following KI-treatment. Permeability after KI treatment (p_KI) was normalized to the pre-treatment value (p_T) as (p_KI-p_T)/p_T. The normalized permeability, along with the biochemical data of kinase target profiles for each of the 12 kinase inhibitors against 291 recombinant protein kinases, served as input to the elastic net regularization algorithm. Model training and feature selection were performed using glmnet package (https://github.com/bbalasub1/glmnet_python, version 2.2.1) implemented in Python 3.7.6. An elastic net mixing parameter of 0.8 was used and a condition-specific cross-validation strategy was employed to identify kinases associated with either barrier disruption or restoration.

### Electron microscopy (SEM and TEM)

BBB microvessels were fixed by perfusing ½ strength Karnovsky’s fixative (2.5% glutaraldehyde, 2% PFA in 0.1 M sodium cacodylate buffer, pH 7.3) through the inlet for 20 min at room temperature, followed by immersion in fresh ½ strength Karnovsky’s fixative for an additional hour. The collagen construct containing 3D microvessels was removed from its acrylic housing device and stored in 6-well plates immersed in fixative at 4°C.

For SEM sample preparation, the bottom collagen layer was peeled to expose open lumen channels for top-down imaging. Samples were rinsed in 0.1 M cacodylate buffer, dehydrated through a graded series of ethanol, and critical point dried (Autosamdri, Tousimis Corp, Rockville, MD). Dried samples were then mounted on stubs, sputter-coated with gold/palladium (Denton Desk IV, Denton Vacuum, Moorestown, NJ), and imaged using a JSM 6610 LV scanning electron microscope at 5 kV (JEOL, Tokyo, Japan).

For TEM sample preparation, the collagen constructs were rinsed in 0.1 M cacodylate buffer, treated with 1% osmium tetroxide for 2 hours at 4°C, rinsed again with cacodylate buffer, dehydrated through a graded series of ethanol and propylene oxide, and embedded in Eponate12 resin (Ted Pella, Inc, Redding, CA). Ultrathin sections (70 nm) were cut using a Leica EM UC7 ultramicrotome, contrasted with uranyl acetate and lead citrate, and imaged on a ThermoFisher Talos L120c transmission electron microscope at 120 kV. Digital images were acquired with a Ceta 16M CMOS 4k x 4k digital camera system.

### Visualization of KiR prediction and EC-permeability analysis

The two-way hierarchically clustered heatmaps were generated using seaborn (https://seaborn.pydata.org, version 0.12.0), a Python data visualization library based on Matplotlib, with Euclidean distance as the clustering metric. The Uniform Manifold Approximation and Projection (UMAP) was performed for dimensionality reduction using the UMAP Python package (https://github.com/lmcinnes/umap) with local neighbors (n_neighbors) of 3, and the results were plotted using Matplotlib (https://matplotlib.org, version 3.5.3) in Python.

### Statistics

All statistical analysis was performed using GraphPad Prism. One-way ANOVA followed by unpaired two-tailed student’s t-test was performed to assess differences in permeability. Statistical significance was defined as a value of p < 0.05. For all imaging and permeability quantification, data was analyzed in a blinded manner. Additional details, including sample sizes and p-values, are provided in the corresponding figure legends.

### Study approval

Collection of TBI plasma samples was approved by the institutional review boards (IRBs) of the participating institutions under study protocols Bloodworks (20141589) and the University of Texas Health Science Center at Houston (HSC-GEN-12-0059).

## Data availability

All datasets generated and analyzed in this study are available from the corresponding authors on reasonable request. The source code and documentation are available through GitHub.

## AUTHOR CONTRIBUTIONS

YZ and AK conceived and supervised the projects. YJS, LW, YZ and AK designed the study. YJS performed all experiments with assistance of LW, CR, SH, NH, KE, JS and JFD. LW performed KIR analysis. YJS, LW, SH, MJ analyzed the data. YJS, LW, AK and YZ wrote the manuscript. All authors edited the manuscripts.

## FUNDING SUPPORT

This work was supported by National Institutes of Health (NIH) grant R61/R33 HL154250 (to A.K., JFD and Y.Z.), R01 AI148802 (to A.K., J.S., Y.Z.), R01 HL152200 (JFD), and R37 CA275954-01A1 (to Y.Z.), Institute for stem cell and regenerative medicine and American Heart Association predoctoral fellowship (to Y.S.).

## ACKNOWLEDGEMENTS

We acknowledge the Lynn and Mike Garvey Imaging Laboratory at the Institute of Stem Cell and Regenerative Medicine. We also thank Steve McFarlene from the Electron Microscope facility at the Fred Hutchinson Cancer Research Institute.

## SUPPLEMENTARY FIGURE LEGENDS

**Supplementary Fig S1. Quantification of barrier permeability using the permeation index.** The permeation index (Δx) quantifies dextran permeation across the endothelium lining the microvessel lumen, and diffusion into the surrounding collagen matrix after 3 min of perfusion with 70 kDa FITC–dextran. A region of interest (ROI) at the vessel midpoint is divided into three windows (ci, ci1, ci2; 100 × 1848 pixels). Pixel intensity profiles are generated for each window, and distances Δxt and Δxb are defined between the 1/3 and 2/3 maximum intensity points (normalized to the local baseline). Δx is the average of Δxt and Δxb across the three windows, normalized to the lumen diameter. Lower Δx indicates reduced dextran spread (intact barrier); higher Δx indicates increased permeability.

Supplementary Fig S2. Time-dependent barrier disruption in 3D BBB microvessels following TBI plasma exposure.

Representative images of 70 kDa dextran perfusion in 3D BBB microvessels 2 and 6 hours after exposure to either pooled normal plasma or plasma from TBI patients. Minimal detectable dextran leakage was observed at 2 hours post-treatment in either condition. Barrier disruption became evident only after 6 hours of TBI plasma incubation. Scale bar: 100 µm.

Supplementary Fig S3. TBI plasma induces heterogeneous increases in vascular permeability.

A. Permeation index before and after 6 h treatment with normal pooled plasma (NPP) show no significant change (N = 4).

B. Permeation index before and after 6-hour treatment with TBI patient plasma show a significant and variable increase in permeability (N = 21). ***P<0.0001; two-tailed student’s t test.

Supplementary Fig S4. Endothelial junctional and morphological changes following thrombin treatment.

A. Quantification of VE-cadherin (VECAD) discontinuity (% of discontinuous junctions) in 3D microvessels under resting conditions, after 15 min thrombin exposure, and after an additional 2h treatment with control DMSO. VECAD remains continuous at baseline, becomes discontinuous after thrombin, and shows further junctional disruption with DMSO.

B-C. Quantification of endothelial cell roundness (aspect ratio) (B) and circularity (C) reveals no immediate morphological change at 15 min post-thrombin, but increased elongation/polarization following 2 h DMSO treatment. ***p < 0.005 and ****p < 0.001 determined by one-way ANOVA and post-hoc Tukey’s test.

**Supplementary Fig S5 Ultrastructural analysis reveals distinct EC barrier morphology in KI-treated 3D BBB microvessels.** (A) SEM imaging of d7 resting BBB microvessels shows (i and ii) a lack of distinct endothelial barrier in resting vessels, indicating a quiescent and intact barrier formation in d7 BBB microvessels. (B) TEM images of resting BBB microvessel shows HBMECs are (i) tethered to the collagen matrix and (ii-iii) form robust adherens and tight junctions (white arrow) with neighboring ECs. Resting vessels also exhibit increased biological activity, demonstrated by active (iv) vesicle transport cycling. (C) SEM imaging of thrombin-activated vessels shows (i) numerous gaps near cell-cell boundaries and (ii) endothelial cell protrusions over the gap formations. (D) TEM images of thrombin-activeated BBB microvessel reveal a similar trend, where endothelial cells are (i) partially delaminated and (ii and iii) show a loss of continuous junctional contact between neighboring ECs. Observations of (iv) lysosomes are noted, but (v) nuclear structure remains normal. (E) SEM imaging of BBB microvessels treated with barrier-restorative KI , Cdk4 inhb, following thrombin activation shows a (i and ii) pronounced junctional boundary, suggesting recovery of cell-cell junctions. (F) TEM images show (i) intact re-endothelialization of ECs along the collagen surface as well as (ii and iii) re-establishment of cell-cell junctions (white arrow). Although (iv) cells do not return to a biologically active state as seen in resting vessels, (v) nuclei remain normal. SEM imaging of (G) BBB microvessels treated with barrier-disruptive KI (Staurosporine) following thrombin activation shows (i and ii) pronounced delamination of endothelial cells and membrane blebbing, suggesting that Staurosporine treatment induces EC apoptosis. (H) TEM images show (i) severe detachment of ECs along the collagen surface as well as (ii) formation of vacuoles and (iii and iv) autophagosomes. (v) Nuclei also exhibit significant condensation.

Supplementary Fig S6. Kinase inhibitors differentially modulate barrier function in 3D BBB microvessels activated with low dose thrombin (0.2 U/mL).

A. Fluorescence images of 70 kDa dextran-perfused BBB microvessels at baseline (D–1), after 15 min of thrombin treatment (0.2 U/mL), and after an additional 2 h of KI treatment. Conditions include DMSO control, barrier-restorative KIs (Cdk2 and Cdk4 inhibitors), and barrier-disruptive KIs (Staurosporine, K-252a, and Dasatinib).

B. Quantification of permeability index normalized to post-thrombin values, enabling assessment of barrier restoration or exacerbation following KI treatment.

Supplementary Fig S7. KIR prediction of barrier-regulatory kinase targets from low-dose thrombin-activated 3D BBB microvessels.

Heatmap of predicted kinase activity based on KiR modeling using permeation index data from 12 kinase inhibitor (KI) treatments in 3D BBB microvessels activated with 0.2 U/mL thrombin. The x-axis shows a subset of KIs predicted to be barrier-restorative (right of vertical axis) or barrier-disruptive (left). The y-axis lists 24 kinase targets identified under these conditions. Red indicates predicted kinase activation; blue indicates inhibition.

Supplementary Fig S8. Predicted KIs from KiR analysis restore barrier function in thrombin-activated 3D BBB microvessels.

A. UMAP visualization of top KiR-predicted barrier-restorative KIs. Most cluster with EGFR/ErbB inhibitors, suggesting shared residual kinase activity profiles.

B. Representative images of 70 kDa FITC-dextran perfusion in BBB microvessels treated with 1 U/mL thrombin followed by selected predicted KIs (p38 inhibitor, MEK1/2 inhibitor, TGF-β inhibitor, and Rapamycin). Top: baseline permeability (D–1); middle: post-thrombin (15 min); bottom: post-KI treatment (2 h).

C. Immunofluorescence images of BBB microvessels fixed immediately after permeability imaging. Samples were stained for VE-cadherin and cytoskeletal markers following 2 h KI treatment after thrombin activation.

**Supplementary Fig S9. Schematic diagram of permeability imaging protocol for TBI patient plasma treated 3D BBB microvessel.** 3D microvessels are treated with TBI plasma for

6hrs followed by 2hr treatment with KIs to validate the effect of KIs for barrier recovery. Permeability imaging was performed after 6hr treatment with TBI patient plasma as well as after 2hr KI treatment within the same microvessels to test the change in barrier permeability.

## SUPPLEMENTARY TABLE LEGENDS

**Supplementary Table S1. 12 original and 6 predicted kinase inhibitors screened in 3D BBB microvessel.**

## Notes

### Competing Interest Statement

The authors have declared no competing interest.

